# Inferring parameters for a lattice-free model of cell migration and proliferation using experimental data

**DOI:** 10.1101/186197

**Authors:** Alexander P. Browning, Scott W. McCue, Rachelle N. Binny, Michael J. Plank, Esha T. Shah, Matthew J. Simpson

## Abstract

Collective cell spreading takes place in spatially continuous environments, yet it is often modelled using discrete lattice-based approaches. Here, we use data from a series of cell proliferation assays, with a prostate cancer cell line, to calibrate a spatially continuous individual based model (IBM) of collective cell migration and proliferation. The IBM explicitly accounts for crowding effects by modifying the rate of movement, direction of movement, and the rate of proliferation by accounting for pair-wise interactions. Taking a Bayesian approach we estimate the free parameters in the IBM using rejection sampling on three separate, independent experimental data sets. Since the posterior distributions for each experiment are similar, we perform simulations with parameters sampled from a new posterior distribution generated by combining the three data sets. To explore the predictive power of the calibrated IBM, we forecast the evolution of a fourth experimental data set. Overall, we show how to calibrate a lattice-free IBM to experimental data, and our work highlights the importance of interactions between individuals. Despite great care taken to distribute cells as uniformly as possible experimentally, we find evidence of significant spatial clustering over short distances, suggesting that standard mean-field models could be inappropriate.

## 1 Introduction

One of the most common *in vitro* cell biology experiments is called a *cell proliferation assay* (Bosco et al., 2015; Bourseguin et al., 2016; Browning et al., 2017). These assays are conducted by placing a monolayer of cells, at low density, on a two-dimensional substrate. Individual cells undergo proliferation and movement events, and the assay is monitored over time as the density of cells in the monolayer increases (Tremel et al., 2009). One approach to interpret a cell proliferation assay is to use a mathematical model (Warne et al. 2017). Calibrating the solution of a mathematical model to data from a cell proliferation assay can provide quantitative insight into the underlying mechanisms, by, for example, estimating the cell proliferation rate (Tremel et al., 2009; Sengers et al., 2007). A standard approach to modelling a cell proliferation assay is to use a mean-field model, which is equivalent to assuming that individuals within the population interact in proportion to the average population density and that there is no spatial structure, such as clustering (Tremel et al., 2009; Sengers et al., 2007; Maini et al., 2004b; Sarapata and de Pillis, 2014; Sherratt and Murray, 1990). More recently, increased computational power has meant that individual based models (IBMs) have been used to directly model the cell-level behaviour (Binny et al., 2016a; Frascoli et al., 2013; Johnston et al., 2014). IBMs are attractive for modelling biological phenomena because they can be used to represent properties of individual agents, such as cells, in the system of interest (Binny et al., 2016a,b; Frascoli et al., 2013; Peirce et al., 2004; Read et al., 2012; Treloar et al., 2013). Typical IBMs use a lattice, meaning that both the position of agents, and the direction of movement, are restricted (Codling et al., 2008). In contrast, lattice-free IBMs are more realistic because they enable agents to move in continuous space, in any direction. However, this extra freedom comes at the cost of higher computational requirements (Plank and Simpson, 2012).

In this work we consider a continuous-space, continuous-time IBM (Binny et al., 2016b). This IBM is well-suited to studying experimental data from a cell proliferation assay with PC-3 prostate cancer cells (Kaighn et al., 1979), as shown in Figure 1(a)-(d). The key mechanisms in the experiments include cell migration and cell proliferation, and we note that there is no cell death in the experiments on the time scale that we consider. Therefore, agents in the IBM are allowed to undergo both proliferation and movement events. Crowding effects that are often observed in two-dimensional cell biology experiments (Cai et al., 2007) are explicitly incorporated into the IBM as the rates of proliferation and movement in the model are inhibited in regions of high agent density. In this study we specifically choose to work with the PC-3 cell line because these cells are known to be highly migratory, mesenchymal cells (Kaighn et al., 1979). This means that cell-to-cell adhesion is minimal for this cell line, and cells tend to migrate as individuals. We prefer to work with a continuous-space, lattice-free IBM as this framework gives us the freedom to identically replicate the initial location of all cells in the experimental data when we specify the initial condition in the IBM. In addition, lattice-free IBMs do not restrict the direction of movement like a lattice-based approach.

**Fig. 1:**
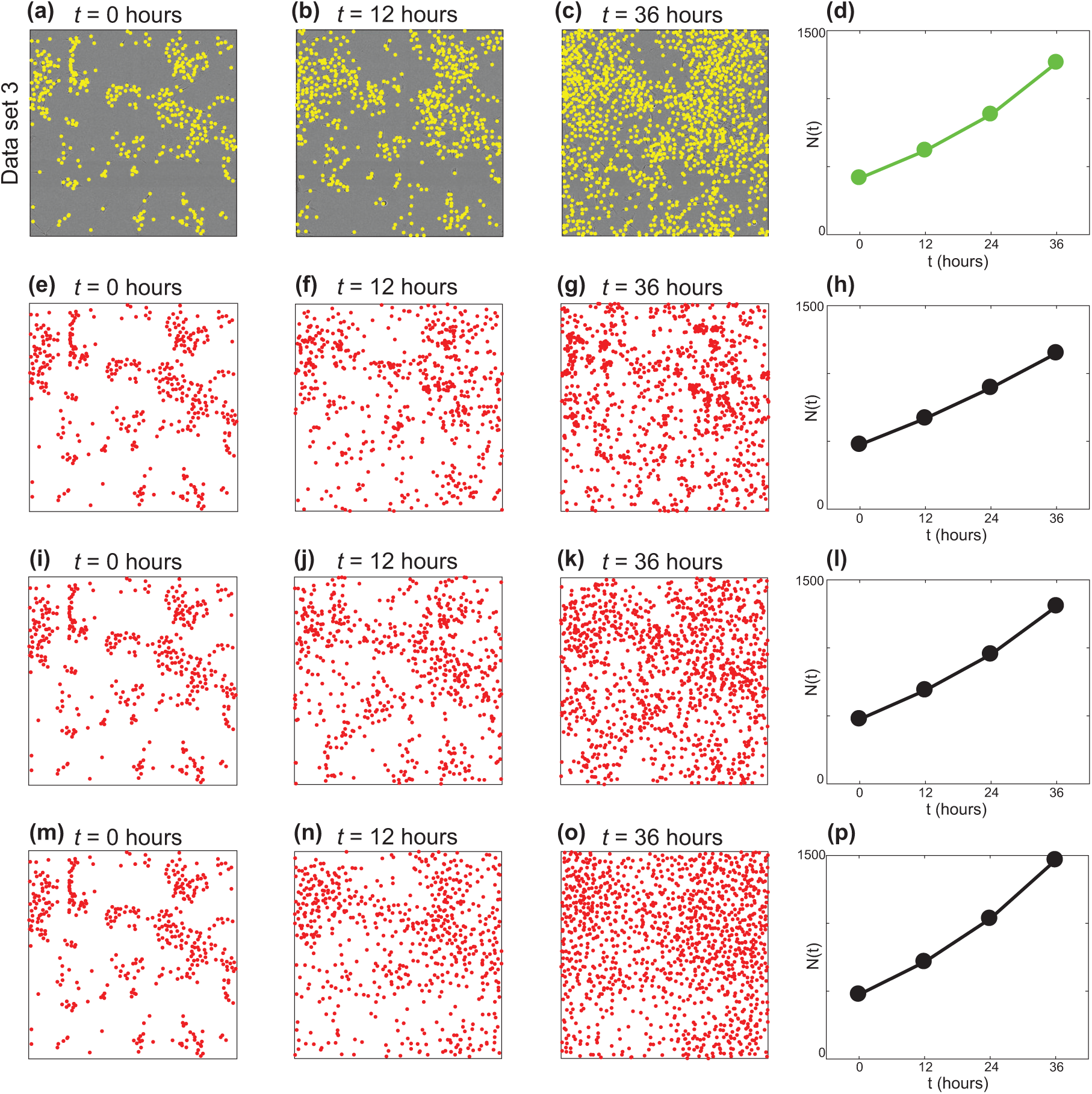
(a)-(c) Experimental data set 3 at *t* = 0, 12 and 36 hours. The position of each cell is identified with a yellow marker. The field of view is a square of length 1440 *µ*m. (d) Population size, *N* (*t*) for experimental data set 3. (e)-(h) One realisation of the IBM with *γ_b_* = 0 *µ*m, leading to an overly clustered distribution of agents. (i)-(l) One realisation of the IBM with *γ_b_* = 6.0 *µ*m, leading to a distribution of agents with similar clustering to the experimental data. (m)-(p) One realisation of the IBM with *γ_b_* = 20 *µ*m, leading to an overly segregated distribution of agents. All IBM simulations are initiated using the same distribution of agents as in (a), with *m* = 1.0 /hour, *p* = 0.040 /hour, and *σ* = 24 *µ*m.

A key contribution of this study is to demonstrate how the IBM can be calibrated to experimental data. In particular, we use approximate Bayesian computation (ABC) to infer the parameters in the IBM. Four sets of experimental images (Supplementary Material 1), each corresponding to an identically-prepared proliferation assay, are considered. The experiments are conducted over a duration of 36 hours, which is unusual because proliferation assays are typically conducted for no more than 24 hours (Browning et al., 2017). Data from the first three sets of experiments (Figure 2) are used to calibrate the IBM and data from the fourth set of images is used to examine the predictive capability of the calibrated IBM. The IBM that we work with was presented very recently (Binny et al., 2016b). The description of the IBM by Binny et al. (2016b) involves a discussion of the mechanisms in the model and the derivation of a spatial moment continuum description (Binny et al., 2016b). Our current work is the first time that experimental data has been used to provide parameter estimates for this new IBM.

**Fig. 2:**
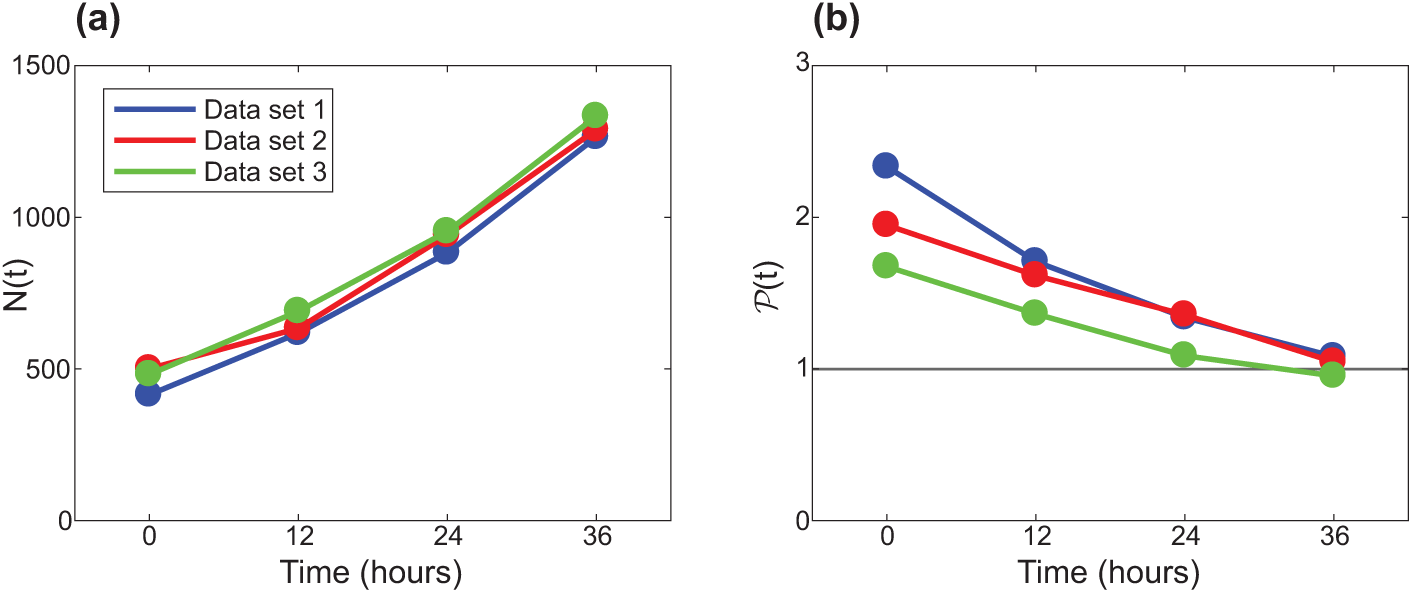
Summary statistics for experimental data sets 1, 2 and 3, shown in blue, red and green, respectively. (a) Population size, *N*(*t*). (b) Local measure of spatial structure, 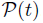, given by Equation 10. Unprocessed experimental data are given in Supplementary Material documents 1 and 2.

Taking a Bayesian approach, we assume that cell proliferation assays are stochastic processes, and model parameters are random variables, allowing us to update information about the model parameters using ABC (Collis et al., 2017; Tanaka et al., 2006). For this purpose we perform a large number of IBM simulations using parameters sampled from a prior distribution. Previous work, based on mean-field models, suggests that the proliferation rate and cell diffusivity for PC-3 cells is *λ ≈* 0.05 /hour and *D ≈* 175 *µ*m^2^/hour, respectively (Johnston et al., 2015). The prior distribution for the IBM parameters are taken to be uniform and to encompass these previous estimates. We generate 10^6^ realisations of the IBM using parameters sampled from the prior distribution, and accept the top 1% of simulations that provide the best match to the experimental data. Our approach to connect the experimental data and the IBM is novel, we are unaware of any previous work that has used ABC to parameterise a lattice-free IBM of a cell proliferation assay. One possible reason why ABC methods are not routinely used to calibrate lattice-free models of cell migration and cell proliferation with crowding effects is because of high computational requirements (Fröhlich et al., 2016). For example, we find that the typical run time to simulate our experiments is approximately 2 seconds on a standard desktop machine using C++. This means that simulating 10^6^ realisations for inference with three unknown parameters becomes challenging. All work presented here is simulated on a High Performance Computing cluster to manage these computational limitations (QUT High Performance Computing, 2017).

Applying the ABC algorithm to data from three sets of identically prepared experiments leads to three similar posterior distributions. This result provides confidence that the IBM is a realistic representation of the cell proliferation assays and leads us to produce a combined posterior distribution from which we use the mean to give point estimates of the model parameters. To provide further validation of the IBM, we use the combined posterior distribution and the IBM to make a prediction of the fourth experimental data set. Simulating the IBM with parameters sampled from the combined posterior distribution allows us to predict both the time evolution of the population size, *N*(*t*), and a measure of the density of pairs of cells, 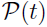, which provides a measure of spatial structure. These results indicate that the *in silico* predictions are consistent with the experimental observations.

This manuscript is organised as follows. Sections 2.1-2.2 describe the experiments and the IBM, respectively. In Section 2.3 we explain how to apply the ABC algorithm to estimate the IBM parameters. In Section 3 we present the marginal posterior distributions of the IBM parameters using data from the first three sets of experiments. The predictive power of the calibrated IBM is demonstrated by using the combined posterior distributions to predict the fourth experimental data set. The predictive power of the calibrated IBM is compared with a stochastic analogue of the standard mean-field logistic equation (Murray, 2002). While both models can accurately predict *N*(*t*), the logistic equation provides no information about the spatial structure in the experimental data. Finally, in Section 4, we conclude and summarise opportunities for further research.

## 2 Material and methods

### 2.1 Experimental methods

We perform a series of proliferation assays using the IncuCyte ZOOM^TM^ live cell imaging system (Essen BioScience, MI USA) (Jin et al., 2017). All experiments are performed using the PC-3 prostate cancer cell line (Kaighn et al., 1979). These cells, originally purchased from American Type Culture Collection (Manassas, VA, USA), are a gift from Lisa Chopin (April, 2016). Cells are propagated in RPMI 1640 medium (Life Technologies, Australia) with 10% foetal calf serum (Sigma-Aldrich, Australia), 100 U/mL penicillin, and 100 *µ*g/mL streptomycin (Life Technologies), in plastic tissue culture flasks (Corning Life Sciences, Asia Pacific). Cells are cultured in 5% CO_2_ and 95% air in a Panasonic incubator (VWR International) at 37 °C. Cells are regularly screened for *Mycoplasma*.

Approximately 8,000 cells are distributed in the wells of the tissue culture plate as uniformly as possible. After seeding, cells are grown overnight to allow for attachment and some subsequent growth. The plate is placed into the IncuCyte ZOOM^TM^ apparatus, and images showing a field of view of size 1440 *×* 1440 *µ*m are recorded every 12 hours for a total duration of 36 hours. Experimental images for experimental data set three is shown in Figure 1(a)-(c). Images from the other three data sets are provided in Supplementary Material 1. ImageJ is used to determine the approximate locations of individual cells in all images, this data is given in Supplementary Material 2. Summary statistics, *N*(*t*) and *P*(*t*), for the first three experimental data sets are given in 2(a)–(b).

### 2.2. Mathematical model

#### 2.2.1. Individual based model

We consider an IBM describing the proliferation and movement of individual cells (Binny et al., 2016a,b Since cell death is not observed in the experiments, the IBM does not include agent death. The IBM allows the net proliferation rate and the net movement rate of agents to depend on the spatial arrangement of other agents. To be consistent with previous experimental observations, the IBM incorporates a biased movement mechanism so that agents tend to move away from nearby crowded regions (Cai et al., 2007). We use the IBM to describe the dynamics of a population of agents on a square domain of length *L* = 1440 *µ*m to match the field-of-view of the experimental data (Figure 1(a)-(c)). Agents in the model are treated as a series of points which we may interpret as a population of uniformly-sized discs with diameter *σ* = 24 *µ*m (Supplementary Material 1). Each agent has location **x**_*n*_ = (*x*_1_*, x*_2_), for *n* = 1*, …, N* (*t*). Since the field-of-view of each image is much smaller than the size of the well in the tissue culture plate, we apply periodic boundary conditions (Jin et. al., 2017).

Proliferation and movement events occur according to a Poisson process over time (Binny et al., 2016b) The *n*th agent is associated with neighbourhood-dependent rates, *P_n_ ≥* 0 and *M_n_ ≥* 0, of proliferation and movement, respectively. These rates consist of intrinsic components, *p >* 0 and *m >* 0, respectively. Crowding effects are introduced by reducing the intrinsic rates by a contribution from other neighbouring agents. These crowding effects are calculated using a kernel, *w*^(*•*)^(*r*), that depends on the separation distance, *r ≥* 0, so that

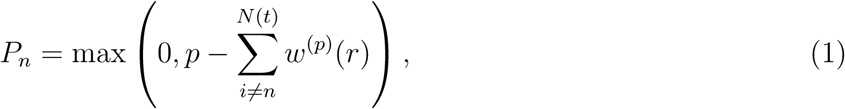

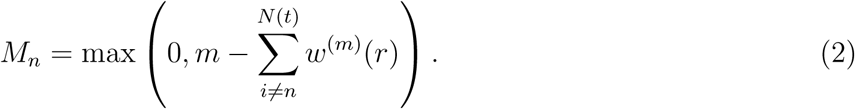

Following Binny et al.,(2016), we specify the kernels to be Gaussian with width corresponding to the cell diameter, *σ*, giving

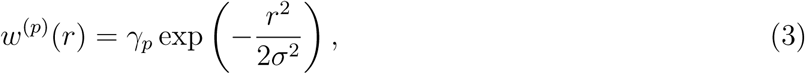

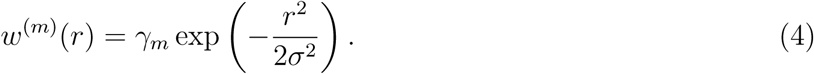

Here, *γ_p_* is the value of *w*^(*p*)^(0) and *γ_m_* is the value of *w*^(*m*)^(0). These parameters provide a measure of the strength of crowding effects on agent proliferation and movement, respectively. The kernels, *w*^(*p*)^(*r*) and *w*^(*m*)^(*r*), ensure that the interactions between pairs of agents separated by more than roughly 2-3 cell diameters lead to a negligible contribution. For computational efficiency, we truncate the Gaussian kernels so that *w*^(*p*)^(*r*) = *w*^(*m*)^(*r*) = 0, for *r ≥* 3*σ* (Law et al., 2003).

To reduce the number of unknown parameters in the IBM, we specify *γ_p_* and *γ_m_* by invoking an assumption about the maximum packing density of the population. Here we suppose that the net proliferation and net movement rates reduce to zero when the agents are packed at the maximum possible density, which is a hexagonal packing (Figure 3(a)). For interactions felt between the nearest neighbours only (Figure 3(b)), we obtain

**Fig. 3:**
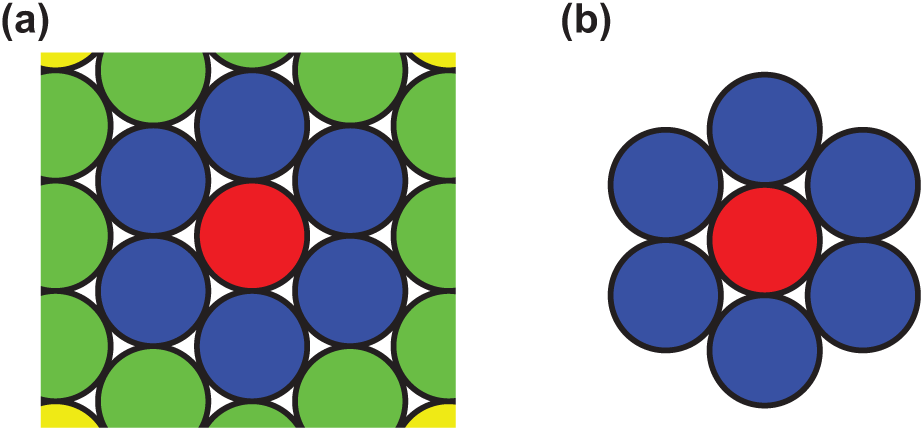
(a) Hexagonal packing of uniformly sized discs. The focal agent (red) is surrounding by six nearest neighbouring agents (blue), and twelve next nearest neighbouring agents (green). (b) Hexagonal packing around a focal agent (red) showing the six nearest neighbours only.

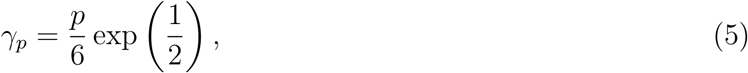

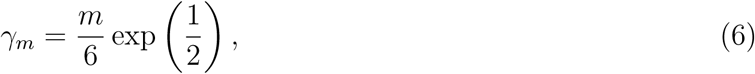

which effectively specifies a relationship between *γ_p_* and *p*, and between *γ_m_* and *m*. Note that this assumption does not preclude a formation of agents in which some pairs have a separation of less than *σ* and densities greater than hexagonal packing, which can occur by chance.

When an agent at **x**_*n*_ proliferates, the location of the daughter agent is selected by sampling from a bivariate normal distribution with mean **x**_*n*_ and variance *σ*^2^ (Binny et al., 2016b). Since mesenchymal cells in two-dimensional cell culture are known to move with a directional movement bias away from regions of high density (Cai et al., 2007), we allow the model to incorporate a bias so that the preferred direction of movement is in the direction of decreasing agent density. For simplicity, the distance that each agent steps is taken to be a constant, equal to the cell diameter, *σ* (Plank and Simpson, 2012).

To choose the movement direction, we use a crowding surface, *B*(**x**), to measure the local crowd-edness at location **x**, given by

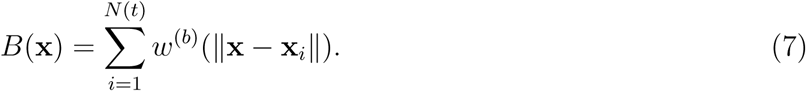

The crowding surface is the sum of contributions from every agent, given by a bias kernel, *w*^(*b*)^(*r*). The contributions depend on the distance between **x** and the location of the *i*th agent, **x**_*i*_, given by *r* = ║**x** *−* **x**_*i*_║. Again, we choose *w*^(*b*)^ to be Gaussian, with width equal to the cell diameter, and repulsive strength, *γ_b_ ≥* 0, so that

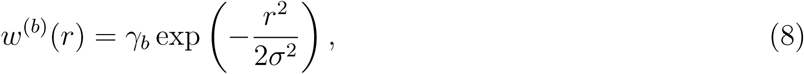

where *γ_b_* is value of *w*^(*b*)^(0), and has dimensions of length. Note that *B*(**x**) is an increasing function of local density, and approaches zero as the local density decreases. A typical crowding surface is shown in Figure 4(b) for the arrangement of agents in Figure 4(a).

**Fig. 4:**
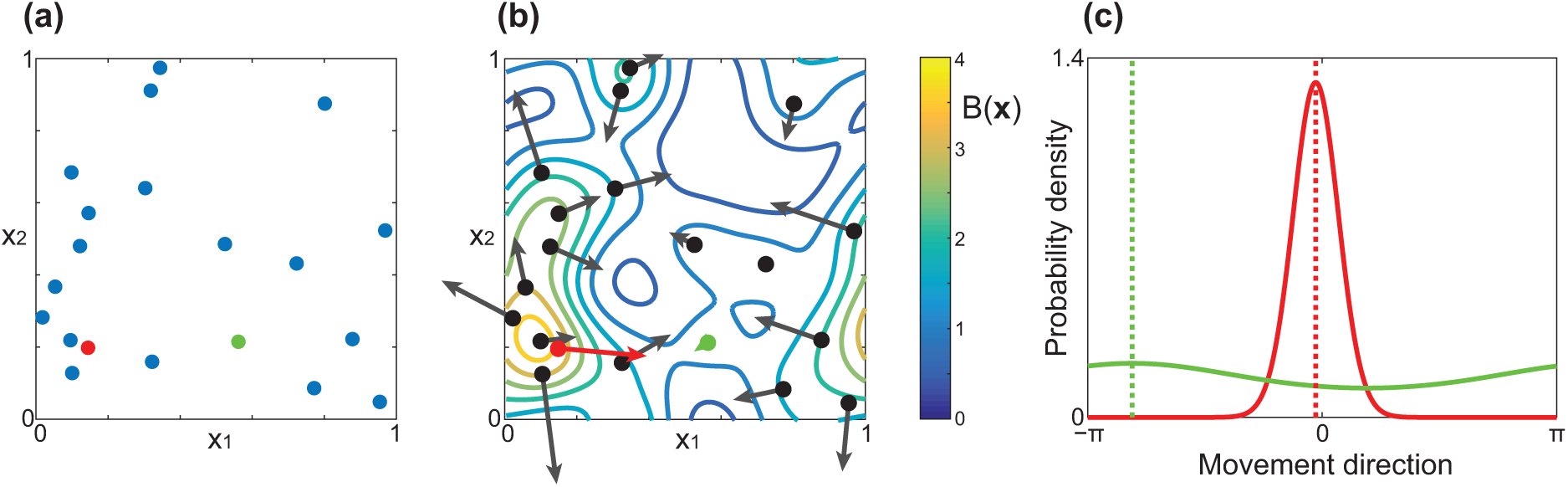
(a) Example distribution of agents on a 1 *×* 1 periodic domain. (b) Level curves of the corresponding crowding surface, *B*(**x**), for this arrangement of agents. The arrows show the preferred direction of movement, **B**_*n*_. To illustrate how the direction of movement is chosen, (c) shows the probability density of the von Mises distribution for the red and green agents highlighted in (a) and (b). The preferred direction, arg(**B**_*n*_), is shown as dotted vertical lines for both agents. The red agent is in a crowded region so ║**B**_*n*_║ is large, meaning that the agent is likely to move in the preferred direction arg(**B**_*n*_). The green agent is in a low density region and **B**_*n*_ is small, meaning that the bias is very weak and the agent’s direction of movement is almost uniformly distributed. To illustrate the effects of the crowding surface as clearly as possible, we set *γ_b_* = 1, *σ* = 0.1, *L* = 1 in this schematic figure to draw attention to the gradient of the crowding surface.

To determine the direction of movement we use the shape of *B*(**x**) to specify the bias, or preferred direction, of agent *n*, **B**_*n*_, given by

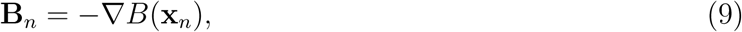

which gives the magnitude and direction of steepest descent. Results in Figure 4(b) show **B**_*n*_ for the arrangement of agents in Figure 4(a). To determine the direction of movement, we consider the magnitude and direction of **B**_*n*_, and sample the actual movement direction from a von Mises distribution, von Mises(arg(**B**_*n*_), ║**B**_*n*_║) (Binny et al., 2016b; Forbes et al., 2011). Therefore, agents are always most likely to move in the direction of **B**_*n*_, however as ║**B**_*n*_║ → 0, the preferred direction becomes uniformly distributed.

To illustrate how the direction of movement is chosen, we show, in Figure 4(b), the bias vector for each agent, **B**_*n*_. Note that **B**_*n*_ does not specify the movement step length, and the direction of **B**_*n*_ does not necessarily specify the actual direction. Rather, arg(**B**_*n*_) specifies the preferred direction. To illustrate this property, we highlight two agents in Figure 4(a). The red agent is located on a relatively steep part of the crowding surface, so ║**B**_*n*_║ is large. The green agent is located on a relatively flat part of the crowding surface, so ║**B**_*n*_║ is close to zero. Figure 4(c) shows the von Mises distributions for the red and green agent. Comparing these movement distributions confirms that the crowded red agent is more likely to move in the direction of **B**_*n*_. The bias is weak for the green agent, so the direction of movement is almost uniformly distributed since ║**B**_*n*_║ is smaller.

IBM simulations are performed using the Gillespie algorithm (Gillespie, 1977). To initialise each simulation we specify the initial number and initial location of agents to match to the experimental images at *t* = 0 hours (Supplementary Material 1) for experimental data sets 1, 2, 3 and 4. In all simulations we set *σ* = 24 *µ*m and *L* = 1440 *µ*m. The remaining three parameters, *m*, *p* and *γ_b_*, are varied with the aim of producing posterior distributions using a Bayesian framework.

If *γ_m_* = *γ_b_* = 0, and the variance of the dispersal distribution is large, the IBM corresponds to logistic growth (Binny et al., 2016b, Browning at al. 2017). Under these simplified conditions, a uniformly distributed initial population of agents will grow, at rate *p*, to eventually reach a uniformly distributed maximum average density of 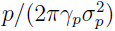. We do not consider this case here as our initial distribution of cells in the experiments is clustered, and so the logistic growth model is, strictly speaking, not valid (Binny et al., 2016b).

#### 2.2.2. Summary statistics

To match the IBM simulations with the experimental data we use properties that are related to the first two spatial moments (Law et al., 2003). The first spatial moment, the average density, is characterised by the number of agents in the population, *N*(*t*). The second spatial moment characterises how agents are spatially distributed, and is often reported in terms of a pair correlation function (Binny et al., 2016a,b; Law et al., 2003). In the Supplementary Material 1 document we present the pair correlation function for all four experimental data sets. These results show that we have a fairly typical pair correlation function that contains, at most, one maximum (Binder and Simpson, 2013). Therefore, instead of using all details contained in the pair correlation function, we use a simplified measure of spatial structure. We consider a local measure of pair density within a distance of *R µ*m, given by

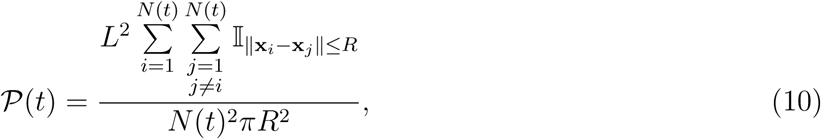

where I is an indicator function so that the double sum in Equation 10 gives twice the number of distinct pairs within a distance *R*. For all results presented in the main document we set *R* = 50 *µ*m. Therefore, 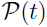 is the ratio of the number of pairs of agents, separated by a distance of less than 50 *µ*m, to the expected number of pairs of agents separated by a distance of less than 50 *µ*m, if the agents were randomly distributed. This means that, 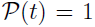 corresponds to randomly placed agents; 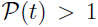 corresponds to a locally clustered distribution; and, 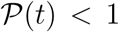 corresponds to a locally segregated distribution.

To ensure that our choice of setting *R* = 50 *µ*m is adequate, we also repeat some results with *R* = 100 *µ*m. This exercise leads to very similar posterior distributions, confirming that working with *R* = 50 *µ*m is sufficient (Supplementary Material 1).

### 2.3. Approximate Bayesian computation

We consider *m, p* and *γ_b_* as random variables, and the uncertainty in these parameters is updated using observed data (Collis et al., 2017; Tanaka et al., 2006). To keep the description of the inference algorithm succinct, we refer to the unknown parameters as **Θ** = *(m, p, γ_b_)*.

In the absence of any experimental observations, information about **Θ** is characterised by specified prior distributions. The prior distributions are chosen to be uniform on an interval that is wide enough to encompass previous estimates of *m* and *p* (Johnston et al., 2015). To characterise the prior for *γ_b_*, we note that this parameter is related to a length scale over which bias interactions are felt. Preliminary results (not shown) use a prior in the interval 0 *≤ γ_b_ ≤* 20 *µ*m and suggest that a narrow prior in the interval 0 *≤ γ_b_ ≤* 10 *µ*m is appropriate. In summary, our prior distributions are uniform and independent, given by

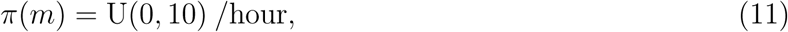

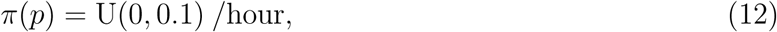

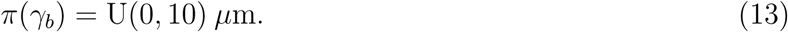

We always summarise data, **X**, with a lower-dimensional summary statistic, *S*. Data and summary statistics from the experimental images are denoted **X**_obs_ and *S*_obs_, respectively. Similarly, data and summary statistics from IBM simulations are denoted **X**_sim_ and *S*_sim_, respectively. Information from the prior is updated by the likelihood of the observations, *π*(*S*_obs_*|***Θ**), to produce posterior distributions, *π*(**Θ***|S*_obs_). We employ the most fundamental ABC algorithm, known as ABC rejection (Liepe et al., 2014; Tanaka et al., 2006), to sample from the approximate posterior distribution. The approximate posterior distributions are denoted *π_u_*(**Θ***|S*_obs_).

In this work we use a summary statistic that is a combination of *N*(*t*) and 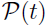 at equally spaced time intervals. A discrepancy measure, *ρ*(*S*_obs_*, S*_sim_), is used to assess the closeness of *S*_obs_ and *S*_sim_,

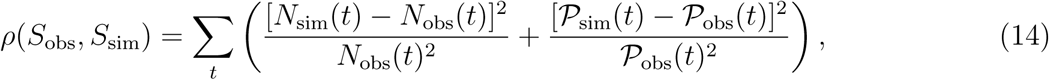

Algorithm 1 is used to obtain 10^6^*u* samples, 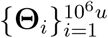, from the approximate joint posterior distribution, *π_u_*(**Θ***|S*_obs_), for each data set. Here, *u* ≪ 1 is the accepted proportion of samples.

To present marginal posterior samples, we use a kernel density estimate to form smooth, approximate marginal posterior distributions, for each parameter, and each data set using the `ksdensity` function in MATLAB (Mathworks, 2017). Point estimates of parameters are always given as the mean of the posterior distribution, and always presented to two significant figures.

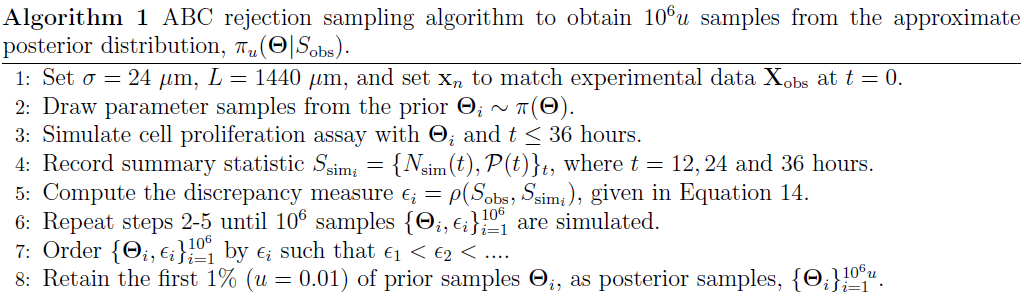

#### 2.3.1. Sampling from the combined posterior distribution

Samples from the posterior distributions for each experimental data set are given in Supplementary Material 1. Kernel density estimates for the marginal posterior distributions for each experimental data set are given in Figure 5. Visually, the posterior distributions for each experimental data set appear to be similar, therefore we are motivated to form a combined posterior distribution, 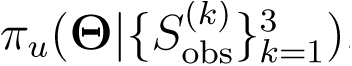, were 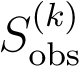 is the summary statistic from the *k*th experimental data set. We use ABC rejection to sample from the combined posterior distribution according to Algorithm 2. That is, Algorithm 2 is designed to sample the combined posterior distribution by retaining the top 1% of parameter combinations that provide the best fit to all three experimental data sets.

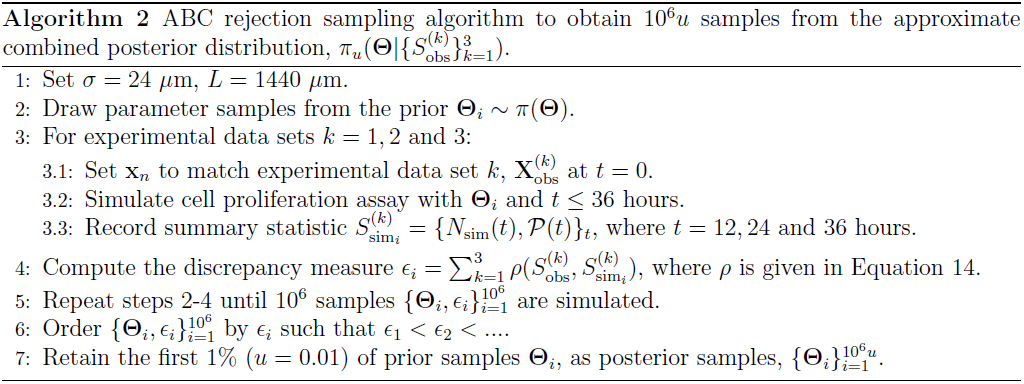

**Fig. 5:**
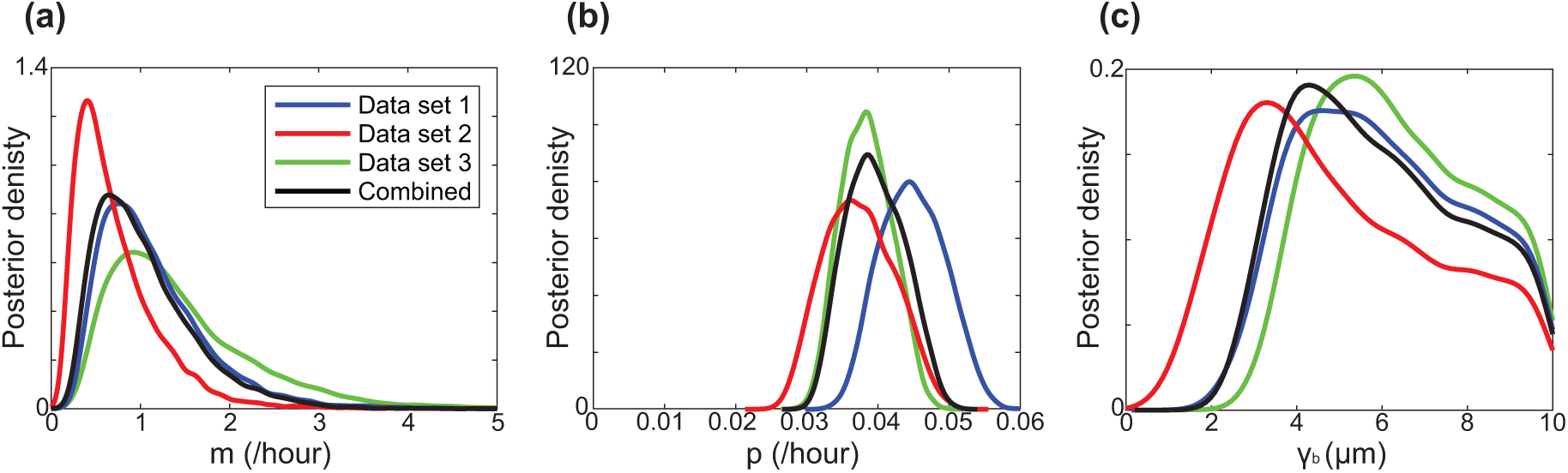
(a)-(c) Kernel-density estimates of the approximate marginal posterior distributions for each data set, for parameters *m*, *p* and *γ_b_*, respectively, with *u* = 0.01. The combined posterior distribution (black) is superimposed. The point estimates from the combined posterior distribution are *m* = 1.0 /hour, *p* = 0.040 /hour and *γ_b_* = 6.0 *µ*m. All distributions are scaled so that the area under the curve is unity.

#### 2.3.2. Predicting experimental data set 4 using the combined posterior distribution

To test the predictive power of the calibrated IBM, we use 10^4^ parameter samples from the combined posterior distribution and simulate the IBM initialised with the actual initial arrangement of cells in data set 4 at *t* = 0. For each parameter combination *S*_sim_ is recorded at 12 hour intervals, and used to construct distributions of *N*(*t*) and 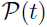. These distributions are represented as box plots and compared with summary statistics from experimental data set 4.

#### 2.3.3. Calibrating the standard mean-field logistic model to the experimental data

To illustrate the importance of considering individual level details in the IBM, we also calibrate the logistic growth model to experimental data sets 1, 2 and 3. The logistic growth model describes the IBM when spatial structure is neglected (Law et al., 2003; Binny et al., 2016b). The logistic growth model is given by

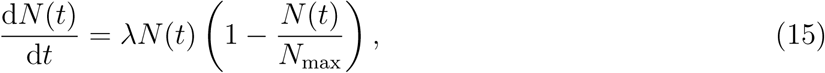

where *λ* is the cell proliferation rate and *N*_max_ is the maximum number of agents. To find estimates of *λ* and *N*_max_ to best match our experimental data we simulate the stochastic logistic model using the Gillespie algorithm (Gillespie, 1977; Fröhlich et. al., 2016). Proliferation events are treated as a Poisson process, with the rate given by the right hand side of Equation 15. Details of the ABC rejection algorithm used to estimate *λ* and *N*_max_ are given in Supplementary Material 1.

## 3 Results and discussion

To qualitatively illustrate the importance of spatial structure we show, in rows 2-4 of Figure 1, snapshots from the IBM with different choices of parameters. In each case the IBM simulations evolve from the initial condition specified in Figure 1(a). Results in the right-most column of Figure 1 compare the evolution of *N*(*t*) and we see that the parameter combination in the second row underestimates *N*(*t*), the parameter combination in the fourth row overestimates *N*(*t*), and the parameter combination in the third row produces a reasonable match to the experimental data. A visual comparison of the spatial arrangement of agents in rows 2-4 of Figure 1 suggests that these different parameter combinations may lead to different spatial structures. This illustration of how the IBM results vary with the choice of parameters motivates us to use ABC rejection to estimate the joint distribution of the parameters. To do this we will use summary statistics from three identically prepared, independent sets of experiments. The summary statistics for these experiments, *N*(*t*) and 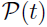, are summarised in Figure 2, and tabulated in Supplementary Material 1.

The approximate marginal posterior distributions for *m*, *p* and *γ_b_* are shown in Figure 5(a)-(c), respectively, for experimental data sets 1, 2 and 3. There are several points of interest to note. In each case, the posterior support is well within the interior of the prior support, suggesting that our choice of priors is appropriate. An interesting feature of the marginal posterior distributions for all parameters is that there is significant overlap for each independent experimental data set. There is some variation in the mean between experimental data sets, for each parameter, which could arise as a consequence of some other variation among experiments, or under the assumption that cell proliferation assays are stochastic processes. The combined marginal posterior distributions are superimposed, and the mean is given by 1.0 /hour, 0.040 /hour and 6.0 *µ*m for *m*, *p* and *γ_b_*, respectively. These point estimates of *p* and *m* give a cell doubling time of ln(2)*/p ≈* 17 hours, and a cell diffusivity of approximately 150 *µ*m^2^/hour, which are typical values for PC-3 cells at low density (Johnston et al., 2015). All results in the main document correspond to retaining the top 1% of samples (*u* = 0.01) and additional results (Supplementary Material 1) confirm that the results are relatively insensitive to this choice.

To assess the predictive power of the calibrated IBM, we attempt to predict the time evolution of a separate, independently collected data set, experimental data set 4, as shown in Figure 6(a)-(d). We use the mean of the combined posterior distribution and the initial arrangement of agents in experimental data set 4 to produce a typical prediction in Figure 6(e)-(h). Visual comparison of the experimental data and the IBM prediction suggests that the IBM predicts a similar number of agents, and a similar spatial structure, with some short range clustering present. To quantify our results, we compare the evolution of *N*(*t*) in Figure 6(i) which reveals an excellent match. Furthermore, we predict the evolution of 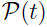 in Figure 6(j) confirming similar trends.

**Fig. 6:**
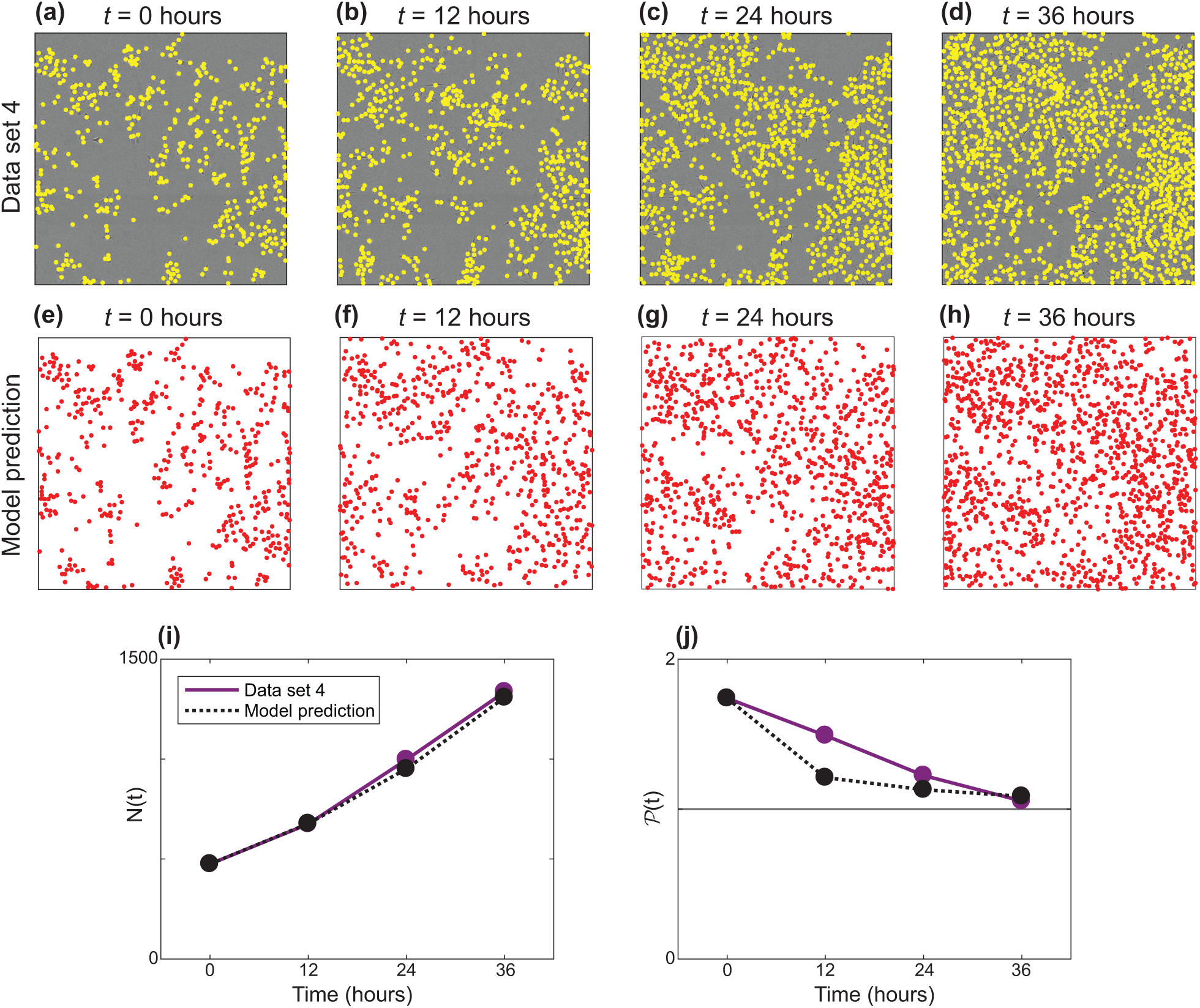
(a)-(d) Experimental images for data set 4. The position of each cell is identified with a yellow marker. The field of view is a square of length 1440 *µ*m. (e)-(h) One realisation of the IBM with parameters corresponding to the posterior mean: *m* = 1.0 /hour, *p* = 0.040 /hour and *γ_b_* = 6.0 *µ*m, with the same initial arrangement of agents as in (a). (i) *N*(*t*) for the experimental data (purple) and the IBM prediction (dashed black). (j) 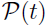 for the experimental data (purple) and the IBM prediction (dashed black). The discrepancy measure (Supplementary Material 1) is given by MSE = 968 for (i), and MSE = 0.0296 for (j).

In addition to examining a single, typical realisation of the calibrated model, we now examine a suite of realisations of the calibrated IBM, and compare results with experimental data set 4. The suite of IBM realisations is obtained by sampling from the joint posterior distribution. Results in Figure 7(a) compare *N*(*t*) from experimental data set 4 with distributions of *N*(*t*) from the suite of IBM simulations, showing an excellent match. The spread of the distributions of *N*(*t*) increases with time, which is expected. Results in Figure 7(b) compare the evolution of 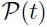 from experimental data set 4 with distributions of 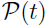 from the suite of IBM simulations, showing the predicted distributions of 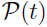 overlap with the experimental data. Overall, the quality of the match between the prediction and the experimental data is high, as the prediction captures both qualitative and quantitative features of the data.

**Fig. 7:**
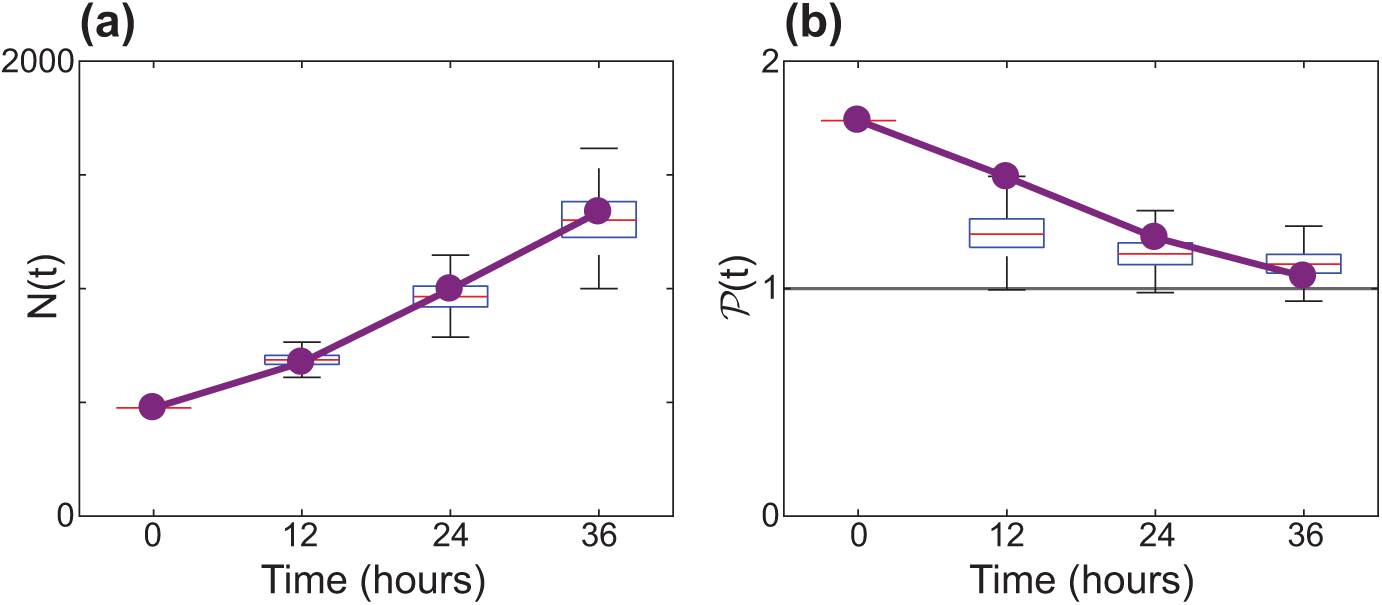
Predictive distributions for *N*(*t*) and 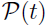, respectively, generated using the IBM. 10^4^ parameter samples are taken from the combined posterior distribution, and a model realisation produced for each sample, initiated as in Figure 6(a). Box plots show the distribution of *N*(*t*) and 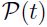 across these realisations in (a) and (b), respectively. The discrepancy measure (Supplementary Material 1) is taken to be between the mean of each boxplot and the observed data, given by MSE = 682 for (a), and MSE = 0.0225 for (b).

We now use ABC rejection to form combined posterior distributions of the parameters in the standard logistic growth model, Equation 15, *λ* and *N*_max_. Results are shown in Figure 8(a)–(b). The point estimates of the combined posterior distributions are *λ* = 0.037 /hour and *N*_max_ = 3600. This estimate leads to a doubling time of approximately 19 hours, which is slightly longer than the doubling time predicted using the calibrated IBM. We then examine a suite of solutions of Equation 15, where we sample from the combined posterior distribution for *λ* and *N*_max_. The predicted distribution of *N*(*t*) is compared with experimental data set 4 in Figure 8(c), revealing an excellent match.

**Fig. 8:**
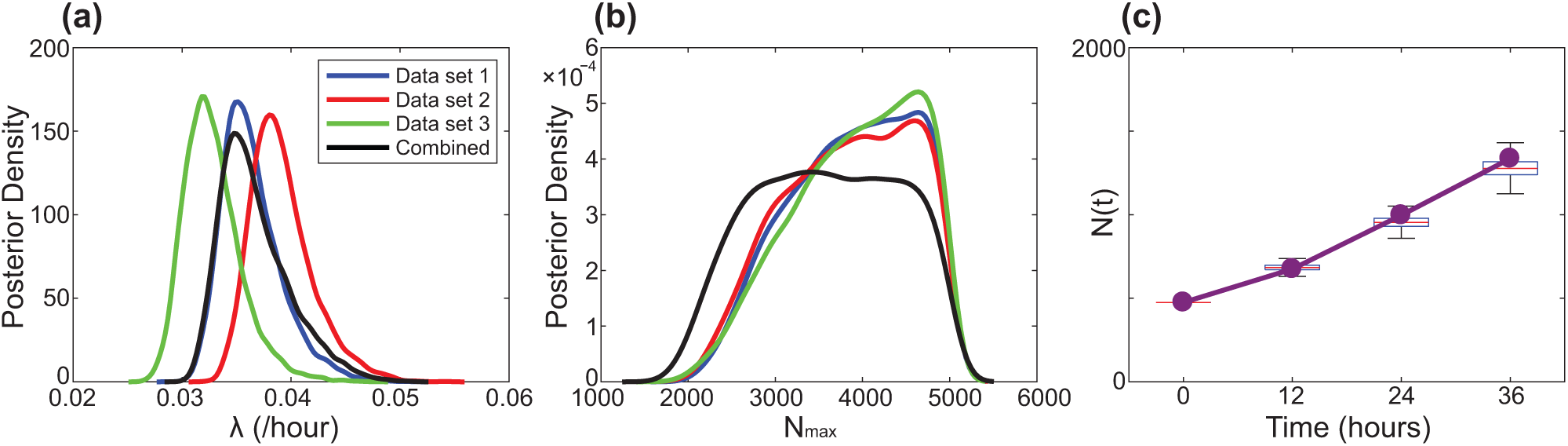
(a)-(b) Kernel-density estimates of the marginal posterior distributions are shown for each data set, for parameters in the stochastic logistic model, Equation 15, *λ* and *N*_max_ in (a) and (b), respectively. The combined posterior distribution (black) is superimposed. The point estimates are *λ* = 0.037 /hour and *N*_max_ = 3600. All marginal distributions are scaled to an area of unity. (c) A predictive distribution for *N*(*t*), generated from the stochastic logistic model, Equation 15. 10^4^ parameter samples are taken from the combined posterior distribution, and a model realisation produced for each sample. Boxplots show the distribution of *N*(*t*) across these realisations. The procedure for sampling from the combined posterior distribution for the stochastic logistic model, and the procedure for solving the stochastic logistic model, are outlined in Supplementary Material 1. The discrepancy measure (Supplementary Material 1) is taken to be between the mean of each boxplot and the observed data, given by MSE = 1668.

Therefore, while both calibrated models provide good predictions for the observed evolution of *N*(*t*), the IBM offers additional insights relating to spatial structure in the cell population, while the logistic model does not provide this level of information. The differences in the way that the logistic model and the IBM treat interactions between individuals could explain why the calibration process leads to different estimates of the proliferation rate. These differences suggest that the interactions between individuals appear to be relevant for our experimental data.

## 4 Conclusions

In this work we explore how to connect a spatially continuous IBM of cell migration and cell proliferation to novel data from a cell proliferation assay. Previous work parameterising IBM models of cell migration and cell proliferation to experimental data using ABC have been restricted to lattice-based IBMs (Johnston et al., 2014). This is partly because ABC methods require large numbers of IBM simulations, and lattice-based IBMs are far less computationally expensive than lattice-free IBMs (Plank and Simpson, 2012). We find it is preferable to work with a lattice-free IBM when dealing with experimental data as a lattice-based IBM requires approximations when mapping the distribution of cells from experimental images to a lattice (Johnston et al., 2014; Johnston et al., 2016). This mapping can be problematic. For example, if multiple cells in an experimental image are equally close to one lattice site, *ad hoc* assumptions have to be introduced about how to arrange those cells on the lattice without any overlap. These issues are circumvented using a lattice-free method.

To help overcome the computational cost of using ABC with a lattice-free IBM, we introduce several realistic simplifying assumptions. The IBM originally presented by Binny et al. (2016b) involves 12 free parameters, which is a relatively large number for standard inference techniques (Schnoerr et al. 2016). The model is simplified by noting that our experiments do not involve cell death, and specifying the width of the interaction kernels to be constant, given by the cell diameter. Another simplification is given by assuming that crowding effects reduce the proliferation and movement rates to zero when the agents are packed at the maximum hexagonal packing density. This leads to a simplified model with three free parameters: *m*, *p* and *γ_b_*. Using ABC rejection, we arrive at posterior distributions for these parameters for three independent experimental data sets. The marginal posterior distributions for the three parameters are similar, leading us to form a combined posterior distribution. The point estimates from the combined posterior distributions for *m* and *p* are consistent with previous parameter estimates (Johnston et al., 2015) and the point estimate for *γ_b_* is consistent with previous observations that mesenchymal cells in this kind of two-dimensional experiment tend to move away from regions of high cell density (Cai et al., 2007).

In the field of mathematical biology, questions about how much detail to include in a mathematical model, and what kind of mathematical model is preferable for understanding a particular biological process are often settled in an *ad hoc* manner, as discussed by Maclaren et al. (2015). Our approach in this work is to use a mathematical model that incorporates just the key mechanisms, with an appropriate number of unknown parameters. Other approaches are possible, such as using much more complicated mathematical models that describe additional mechanisms such as: (i) detailed information about the cell cycle in individual cells (Fletcher et al., 2012); (ii) concepts of leader and follower cells (Kabla, 2012); (iii) explicitly coupling cell migration and cell proliferation to the availability of nutrients and growth factors (Tang at al., 2014); or (iv) including mechanical forces between cells (Stichel at al., 2017). However, we do not include these kinds of detailed mechanisms because our experimental data does not suggest that these mechanisms are relevant to our situation. Furthermore, it is not always clear that using a more complicated mathematical model, with additional mechanisms and additional unknown parameters, necessarily leads to improved biological insight. In fact, simply incorporating additional mechanisms and parameters into the mathematical model often leads to a situation where multiple parameter combinations lead to equivalent predictions which limits the usefulness of the mathematical model (Simpson et al., 2006). In this study, our approach is to be guided by experimental data and our ability to infer the parameters in a mathematical model based on realistic amounts of experimental data (Maclaren et al. 2015). In particular we use three experimental data sets to calibrate the IBM, and an additional data set to separately examine the predictive capability of the calibrated IBM. We find that the process of calibrating the IBM leads to well defined posterior distributions of the model parameters, and that the calibrated IBM produces a reasonable match to the experimental data. The process of calibrating the IBM, and then separately testing the predictive capability of the calibrated IBM, provides some confidence that the level of model complexity is appropriate for our purposes.

An interesting feature of all experimental data at early time, when the cell density is relatively low, is that 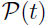 suggests that the cells are clustered at short intervals, and that this clustering becomes less pronounced with time. This observation is very different to the way that previous theoretical studies have viewed the role of spatial structure. For example, previous simulationbased studies assume that some initial random spatial arrangement of cells can lead to clustering at later times (Baker and Simpson, 2010). In contrast, our experimental data suggests it could be more realistic to consider that the spatial structure is imposed by the initial arrangement of cells. Moreover, since all of our experimental data involves some degree of spatial clustering, our work highlights the importance of using appropriate models to provide a realistic representation of key phenomena. Almost all continuum models of collective behaviour in cell populations take the form of ordinary differential equations and partial differential equations that implicitly invoke a mean-field assumption (Tremel et al., 2009; Sengers et al., 2007; Maini et al., 2004b; Sarapata and de Pillis, 2014; Sherratt and Murray, 1990). Such assumptions ignore the role of spatial structure. While pairwise models that avoid mean-field assumptions are routine in some fields, such as disease spreading (Sharkey et al., 2006; Sharkey, 2008) and ecology (Law et al., 2003), models that explicitly account for spatial structure are far less common for collective cell behaviour.

Using our parameter estimates, the continuum spatial moment description could be used to interpret experimental data sets with larger numbers of cells (Binny et al., 2016b), such as experimental images showing a wider field-of-view, or experiments initiated with a higher density of cells. Our approach to estimate the parameters in the model is to work with the IBM since this allows us more flexibility in connecting with the experimental data, such as choosing the initial locations of the agents in the IBM to precisely match the initial locations of cells in the experimental images.

There are many ways that our study could be extended. For example, here we choose a summary statistic encoding information about the first two spatial moments. However, other summary statistics may provide different insight, and it could be of interest to explore the effect of this choice. For example, here we describe the spatial structure over a relatively short interval, approximately 2*σ*. It could be of interest to repeat our analysis using the entire pair correlation function, accounting for spatial structure at all distances. However, here we take a simpler approach and we provide evidence that our results are insensitive to our measure of spatial structure as we obtain similar results when we consider spatial structure over larger distances. Another limitation of our work is that the IBM, which explicitly accounts for interactions between agents, can be computationally expensive to simulate. This limitation can be particularly problematic for computational inference and severely limits the number of parameters that can be dealt with by taking a purely individual-based approach. One promising way of overcoming this difficulty is to make use of more theoretical ways to treat interactions between individuals in an IBM, and to perform inference using a stochastic continuum description, such as the recent work by Schnoerr et al. (2016; 2017).

## 5 Acknowledgements

This work is supported by the Australian Research Council (DP140100249, DP170100474) and the Royal Society of New Zealand Marsden Fund (11-UOC-005). Computational resources provided by the High Performance Computing and Research Support Group at QUT are appreciated. We thank David Warne for technical advice. We also thank the two anonymous referees for their helpful comments.

